# Beyond the vestibulo-ocular reflex: Vestibular input is processed centrally to achieve visual stability

**DOI:** 10.1101/260299

**Authors:** Edwin S. Dalmaijer

## Abstract

The current study presents a re-analysis of data from Zink et al. (1998, *Electroencephalography and Clinical Neurophysiology, 107*), who administered galvanic vestibular stimulation through unipolar direct current. They placed electrodes on each mastoid, and applied both right and left anodal stimulation. Ocular torsion and visual tilt were measured under different stimulation intensities. New modelling introduced here demonstrates that directly proportional linear models fit reasonably well to the relationship between vestibular input and visual tilt, but not to that between vestibular input and ocular torsion. Instead, an exponential model characterised by a decreasing slope and an asymptote fitted best. These results demonstrate that in the results presented by Zink et al., ocular torsion could not completely account for visual tilt. This suggests that vestibular input is processed centrally to stabilise vision when ocular torsion is insufficient. Potential mechanisms and seemingly conflicting literature are discussed.

## Introduction

During everyday movements like walking, the human head and eyes continuously move, yet humans have a relatively stable visual perception of the world. This is due to the vestibulo-ocular reflex. When the vestibular system senses a head movement, it signals directly to the eye muscles, and a compensatory eye movement is produced to realign the visual world (**Figure 1A-C**). Specifically, when the human head makes a rolling movement, a compensatory torsional eye movement in the opposite direction is generated, thereby keeping the visual field stable.

**Figure 1.**
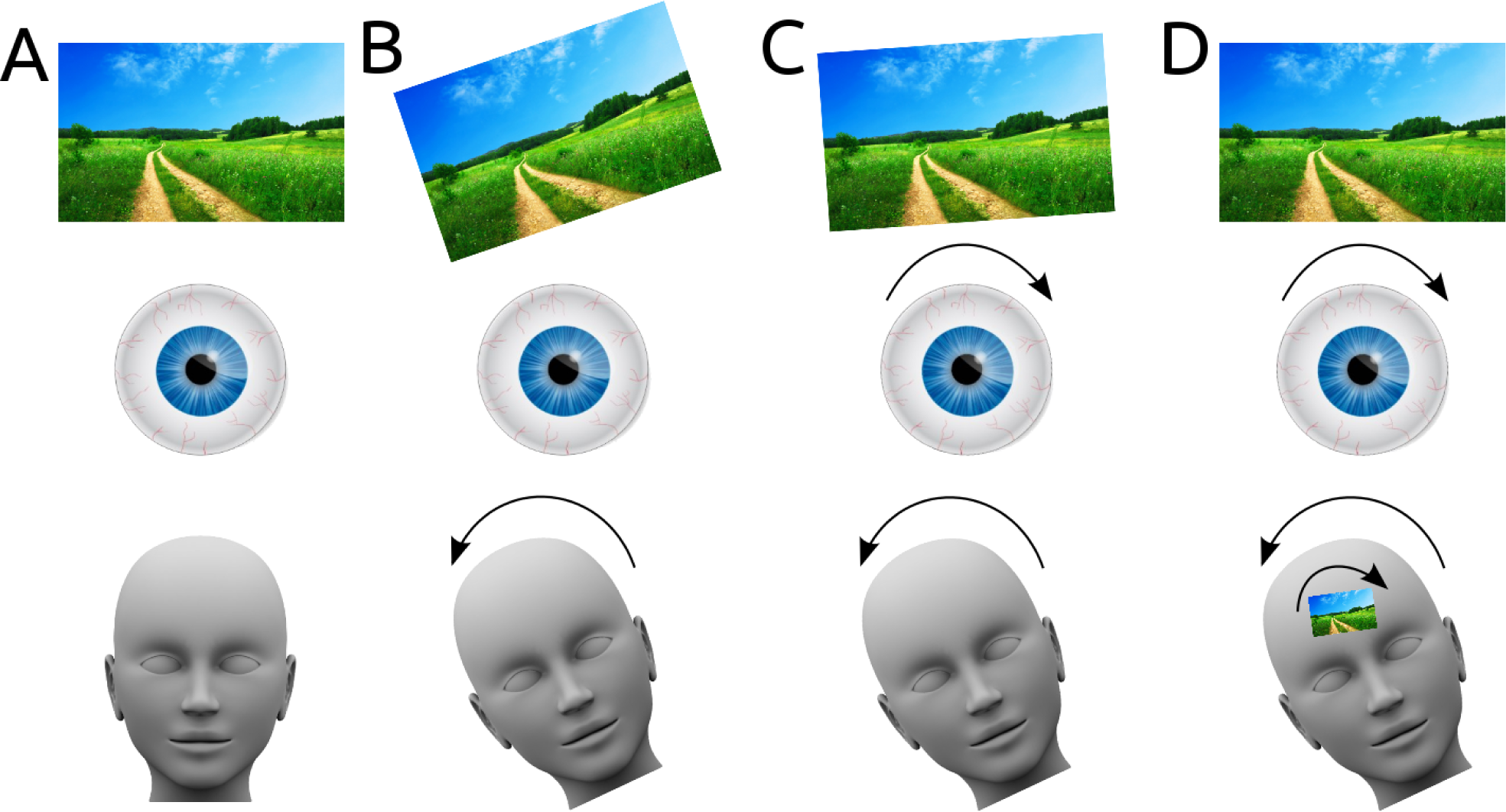
- A) When the head is in an upright position, so are the eyes and the visual world. B) If the eyes were to remain upright with respect to a tilting head, the visual world would tilt. C) Due to the vestibulo-ocular reflex, the eyes rotate in the opposite direction of the rolling head. This counteracts most of the rotation of the visual field. D) Some authors have argued that vestibular input is also processed centrally, to directly tilt visual fields.

However, torsional eye movements induced by the vestibulo-ocular reflex can be prevented by fixating the eyeballs of anaesthetised cats to a metal ring. In these cats, the receptive fields of a proportion of neurons in visual cortex tilted when the cat was tilted, compared to when it was in an upright position (Denney & Adorjanti, 1972; Horn & Hill, 1969; Horn, Stechler, & Hill, 1972). These results suggest that another mechanism might exist to centrally process vestibular information when the vestibulo-ocular reflex is disrupted (**Figure 1D**).

This suggestion is not without controversy: Receptive fields did not tilt as a function of bodily tilt in cats that were not anaesthetised and did not have their eyeballs are not fixated (Schwartzkroin, 1972). Importantly, receptive field tilt *did* occur, but it was uncorrelated with bodily tilt. Schwartzkroin thus suggested that the tilting of receptive fields could have been an effect of non-specific arousal (with the tilting as an arousing stimulus), or could be interpreted as “broadening of direction preference". Therefore, it remains unclear whether central processing of vestibular input influences visual stability beyond its role in the vestibulo-ocular reflex.

Ideally, the hypothesised central processing of vestibular information could be directly tested by comparing the effects of vestibular input on visual tilt and on ocular torsion. If the vestibulo-ocular reflex is sufficient for stabilising the visual world, the ocular torsion induced by vestibular input should be linearly related to the induced visual tilt. Such a study exists: Zink and colleagues reported that both ocular torsion and visual tilt increase with vestibular input (Zink, Bucher, Weiss, Brandt, & Dieterich, 1998).

Human research participants are nor normally have their eyes fixated to metal rings while their body is tilted. Instead, Zink and colleagues induced vestibular input by means of galvanic vestibular stimulation. This technique that has been known for over a hundred years (Day, 1999), with the first reports dating back to around 1900 (Buys, 1909; Hitzig, 1898). Galvanic vestibular stimulation is an electric current (usually applied via both mastoids) that stimulates the vestibular neuronal afferents, and is known to induce spontaneous nystagmus (at intensities over 3 mA) and ocular torsion (Kleine, Guldin, & Clarke, 1999; Schneider, Glasauer, & Dieterich, 2000; Zink et al., 1998).

Eye movements induced by galvanic vestibular stimulation occur due to electrical stimulation of semicircular canal afferents (Schneider, Glasauer, & Dieterich, 2002). Although otolith activation occurs at higher stimulation intensity, its contribution to eye movement varies between individuals (Kleine et al., 1999; Zink et al., 1998). A contemporary model on the contribution of semicircular canal and otolith activation is provided by Day and colleagues (Day, Ramsay, Welgampola, & Fitzpatrick, 2011), who built on earlier models by the same group (Fitzpatrick & Day, 2004) and others (Schneider et al., 2000).

In the aforementioned study by Zink and colleagues, it was reported that both ocular torsion and visual tilt increase with galvanic vestibular stimulation at higher current intensities (Zink et al., 1998). However, induced ocular torsion will invoke visual tilt merely due to the rotation of both eyeballs, but from the analysis by Zink and colleagues it is unclear whether ocular torsion could account for all visual tilt, or whether part of the visual tilt could have been ascribed to central processing of vestibular input.

The current study aims to investigate the relationship between ocular torsion and visual tilt re-analysing the results of Zink and colleagues by modelling the contributions of vestibular stimulation on ocular torsion and visual tilt. Specifically, the models introduced here take into account a hypothesised limit on ocular torsion. If such a limit exists, visual tilt will increasingly depend on central processing when vestibular input increases. Alternatively, if both ocular torsion and visual tilt are linearly related to vestibular input, it would suggest the vestibulo-ocular reflex is sufficient, and no central processing of vestibular input is required to stabilise vision.

## Methods

### Zink et al. (1998)

Relevant data on ocular torsion and visual tilt was extracted from Zink and colleagues (Zink et al., 1998), who conveniently provided it in tables with descriptive statistics of all necessary measures. They used electrodes taped to both mastoids to deliver a unipolar direct current. During a trial, the polarity and intensity of the stimulation was kept constant, but they could be varied between trials. Specifically, Zink and colleagues tested both left-anodal and right-anodal stimulation, and they applied current intensities between 1 and 7 mA.

Stimulation trials lasted for 5 seconds, during which static ocular torsion was measured using a laser-scanning opthalmoscope that recorded the fundus in both eyes on video.

To measure visual tilt, participants were positioned in front of a half-open dome with a diameter of 60 cm that completely covered their visual field, and that was covered with a pattern of randomly placed colour dots. The dome prevented participants from using straight lines in the environment as references. Perceived visual tilt was measured using a centrally presented line that participants could adjust to the level of rotation that they perceived during galvanic vestibular stimulation.

Ocular torsion occurred towards the anode (counter-clockwise under left anodal stimulation, clockwise in right anodal stimulation). Visual tilt occurred away from the anode (clockwise in left anodal stimulation, and counter-clockwise in right anodal stimulation). The results from Zink and colleagues are reproduced in **Table 1**.

**Table 1.** Results from Zink et al. (1998), Electroencephalography and clinical Neurophysiology, 107, p. 200-205. Zink and colleagues applied unipolar direct current with the anode on the right or left mastoid. They measured ocular torsion and visual tilt in degrees of rotation at different galvanic vestibular stimulation intensities. Results reported by Zink and colleagues (and reprinted here) are the average rotation (unsigned), the standard deviation (between round brackets), the minimum and maximum measured values (between square brackets), and the number of participants tested in a particular cell. Ocular torsion occurred towards the anode, whereas visual tilt occurred away from the anode.

| Current strength<br>(mA) | Left anodal stimulation |  | Right anodal stimulation |  |
| --- | --- | --- | --- | --- |
|  | Ocular torsion | Visual tilt | Ocular torsion | Visual tilt |
| **1.0** | 1.0 (0.4)<br>[0.5 – 1.5] N=6 |  | 1.2 (0.3)<br>[0.6 – 1.4] N=6 |  |
| **1.5** | 1.3 (0.1)<br>[1.2 – 1.4] N=2 | 2.2 (0.9)<br>[1.3 – 3.3] N=4 | 1.4 (0.1)<br>[1.3 – 1.4] N=2 | 1.7 (0.5)<br>[1.3 – 2.3] N=4 |
| **2.0** | 2.0 (0.5)<br>[1.3 – 2.5] N=7 | 2.6 (1.4)<br>[1.3 – 6.3] N=12 | 2.1 (0.5)<br>[1.5 – 2.8] N=7 | 2.6 (1.2)<br>[1.0 – 5.8] N=12 |
| **2.5** |  | 3.2 (2.3)<br>[1.2 – 9.4] N=12 |  | 3.1 (2.0)<br>[1.0 – 8.5] N=12 |
| **3.0** | 2.5 (0.8)<br>[1.4 – 3.5] N=7 | 4.9 (1.5)<br>[3.0 – 6.5] N=4 | 3.0 (0.6)<br>[2.2 – 3.5] N=7 | 4.8 (1.8)<br>[2.6 – 6.4] N=4 |
| **4.0** | 2.9 (1.0)<br>[1.2 – 3.8] N=6 |  | 3.3 (1.3)<br>[1.3 – 4.2] N=6 |  |
| **5.0** | 3.2 (1.1)<br>[2.0 – 4.1] N=3 |  | 3.6 (1.9)<br>[1.5 – 4.3] N=3 |  |
| **6.0** | 3.6 (1.3)<br>[2.2 – 4.5] N=3 |  | 4.1 (1.7)<br>[2.2 – 5.2] N=3 |  |
| **7.0** | 3.9 (1.8)<br>[2.6 – 5.2] |  | 4.0 (2.1)<br>[2.5 – 5.4] |  |

For all analyses presented here, the left and right anodal stimulation was averaged within each stimulation intensity.

### Linear models

Zink and colleagues fit linear models that describe both ocular torsion and visual tilt as a function of stimulation intensity (**Equations 1 and 2**). Free variables *a* and *b* in these equations determine the slope and intercept of the function. It should also be noted that the *a* and *b* parameters in the ocular torsion equation are independent from those in the visual tilt equation.

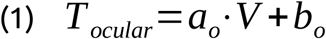

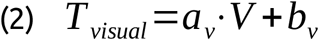

Where *T*_*ocular*_ is ocular torsion in degrees, *V*is vestibular input in mA; *a*_*o*_ is the slope in degrees per mA, and *b*_*o*_ the intercept in degrees in the relationship between ocular torsion and visual tilt. *T*_*visual*_ is visual tilt in degrees; *a*_*v*_ is the slope in degrees per mA, and *b*_*v*_ the intercept in degrees.

### Directly proportional linear models

By accounting for an intercept in **Equations 1**, Zink and colleagues allowed the baseline ocular torsion to be different from 0 degrees at 0 mA of stimulation. Because stimulation can be applied in two directions, according to **Equation 1** the eye is in a different baseline position depending on whether −0 or 0 mA of stimulation is applied. The same is true for **Equation 2** and visual tilt.

Evidently, this cannot be true: At 0 mA of stimulation, the eye and visual field should be un-rotated. The best way to account for this, is by requiring that vestibular input and ocular torsion (or visual tilt) are directly proportional. This idea was implemented in **Equations 3 and 4**, which both have only one free variable that determines the slope of the function.

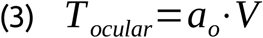

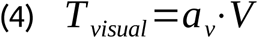

Where *T*_*ocular*_ is ocular torsion in degrees, *V*is vestibular input in mA, and *a*_*o*_ is the slope in degrees per mA in the relationship between ocular torsion and visual tilt. *T*_*visual*_ is visual tilt in degrees, and *a*_*v*_ is the slope in degrees per mA in the relationship between visual tilt and vestibular input.

### Exponential model of ocular torsion

If ocular torsion is indeed limited by an upper bound, neither linear model described above would describe the relationship between galvanic vestibular stimulation and ocular torsion accurately. Instead, as vestibular input increases, the slope of the increase in ocular torsion is expected to decrease, and to become 0 when an asymptote is reached. **Equation 5** described such a relationship, describing that at high levels of vestibular input, ocular torsion will be no higher than asymptote b.

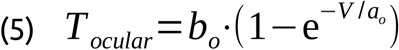

Where *T*_*ocular*_ is the ocular torsion in degrees, *V*is the vestibular input in mA; *a*_*o*_ determines the slope of the function (with lower numbers reflecting a steeper slope), and *b*_*o*_ determines the asymptote of the function (preventing *T*_*ocular*_ to ever rise above *b*_*o*_, regardless of the value of *V*).

### Exponential model of visual tilt

If ocular torsion is indeed limited by an upper bound, visual tilt would increasingly depend on central processing of vestibular stimulation. Thus, as galvanic vestibular stimulation increases, one could expect visual tilt to non-linearly increase. **Equation 6** formalises this relationship as an exponential increase governed by two free variables.

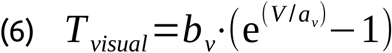

Where *T*_*visual*_ is the visual tilt in degrees, *V* is the vestibular input in mA; *a*_*v*_ determines the slope of the function (with lower numbers reflecting a steeper slope), and *b*_*v*_ determining the linear increase in slope (with higher numbers resulting in steeper slopes).

### Curve fitting

To fit **Equations 1–6** to the data reported by Zink and colleagues (reproduced in **Table 1**), the unsigned results from left-anodal and right-anodal stimulation were first averaged. Then, using least squares estimation in a full exploration of parameter space, the optimal combination of parameter values was assessed for the three types of models outlined above (linear, directly proportional linear, and exponential) for ocular torsion and visual tilt independently.

Fitting was performed in custom Python scripts (Dalmaijer, 2017; Van Rossum & Drake, 2011), using NumPy (Oliphant, 2007) for computations, and Matplotlib (Hunter, 2007) for plotting. This code is available from GitHub on https://github.com/esdalmaiier/zink et al 1998 re-analysis.

## Results

### Parameter estimates

Parameter space was explored within the range 0 - 3.5 with grid resolution of 0.0005 for parameters *a* and *b* in linear models (**Equations 1–4**). The space was explored within the range 0 - 7 with grid resolution 0.001 for parameters *a* and *b* in exponential models (**Equations 5 and 6**).

Parameter estimates are visualised in **Figure 2**, with the best fitting combination of parameters indicated with a circle. The best fits for ocular torsion were *a*_*o*_=0.483 and *b*_*o*_=0.913 for the linear model (**Equation 1**), *a_*o*_*=0.672 for the directly proportional linear model (**Equation 3**), and *a*_*o*_=3.722 and *b*_*o*_=4.714 for the exponential model (**Equation 5**). The best fits for visual tilt were *a*_*v*_=1.421 and *b*_*v*_*=0* for the linear model (**Equation 2**), *a*_*v*_=1.421 in the directly proportional linear model (**Equation 4**), and *a*_*v*_ =2.852 and *b*_*v*_ =2.510 for the exponential model (**Equation 6**).

**Figure 2.**
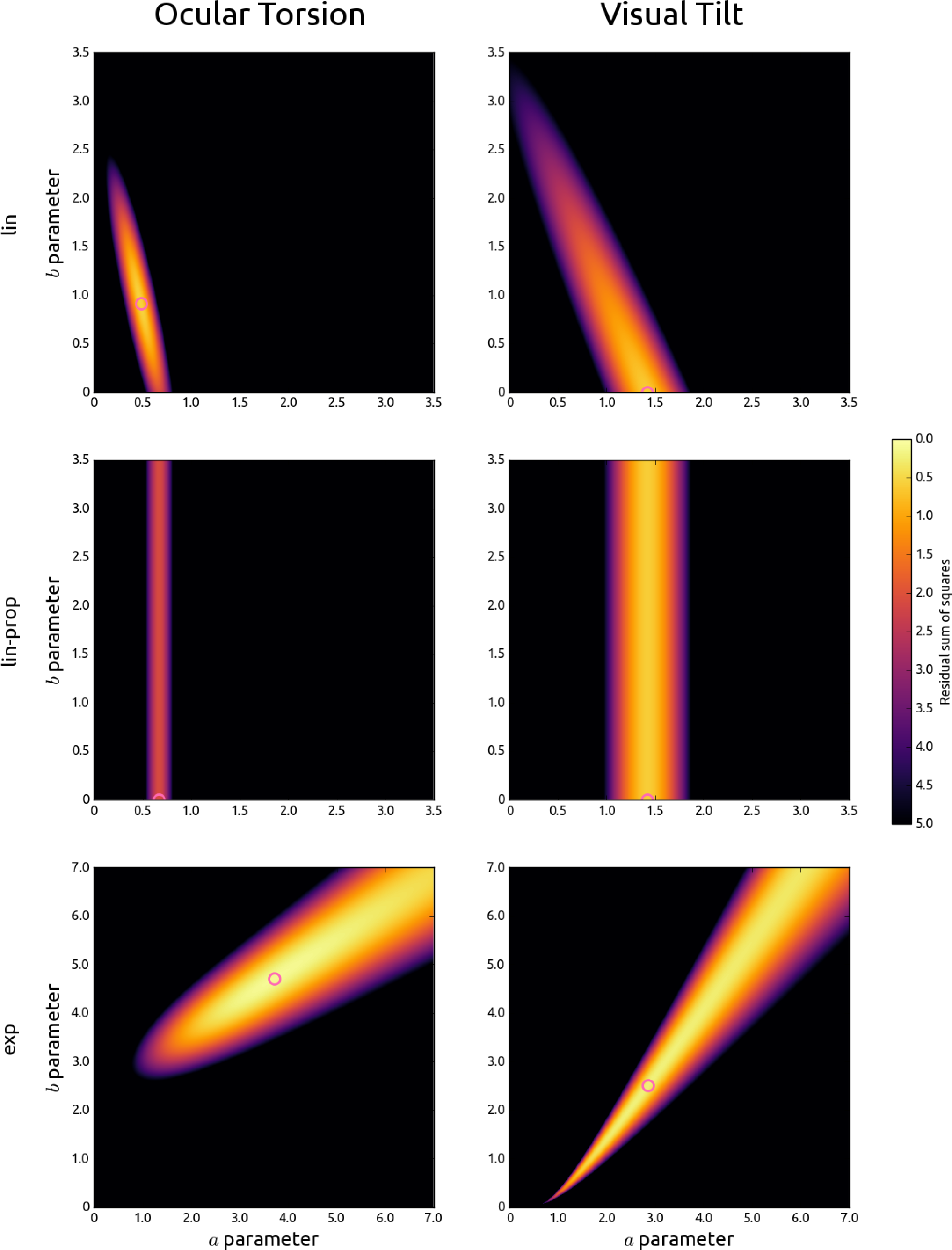
- Visualisation of the residual sum of squares in parameter space as a function of the a (x-axis) and b parameter (y-axis) in linear models (Equations 1 and 2; top row, titled ‘lin), directly proportional linear models (Equations 3 and 4; middle row, titled lin-prop), and exponential models (Equations 5 and 6; bottom row, titled ’exp) of the relationship between galvanic vestibular stimulation and ocular torsion (left column) or visual tilt (right column). Lower values indicate better fits and are indicated by lighter colours. The best fit is indicated by a pink circle.

### Ocular torsion

A directly proportional linear model (**Equation 3**) explained 75 percent of the variance in ocular torsion under galvanic vestibular stimulation (**Figure 3**, in blue). By contrast, an exponential model (**Equation 5**) explained 99 percent of the variance.

**Figure 3.**
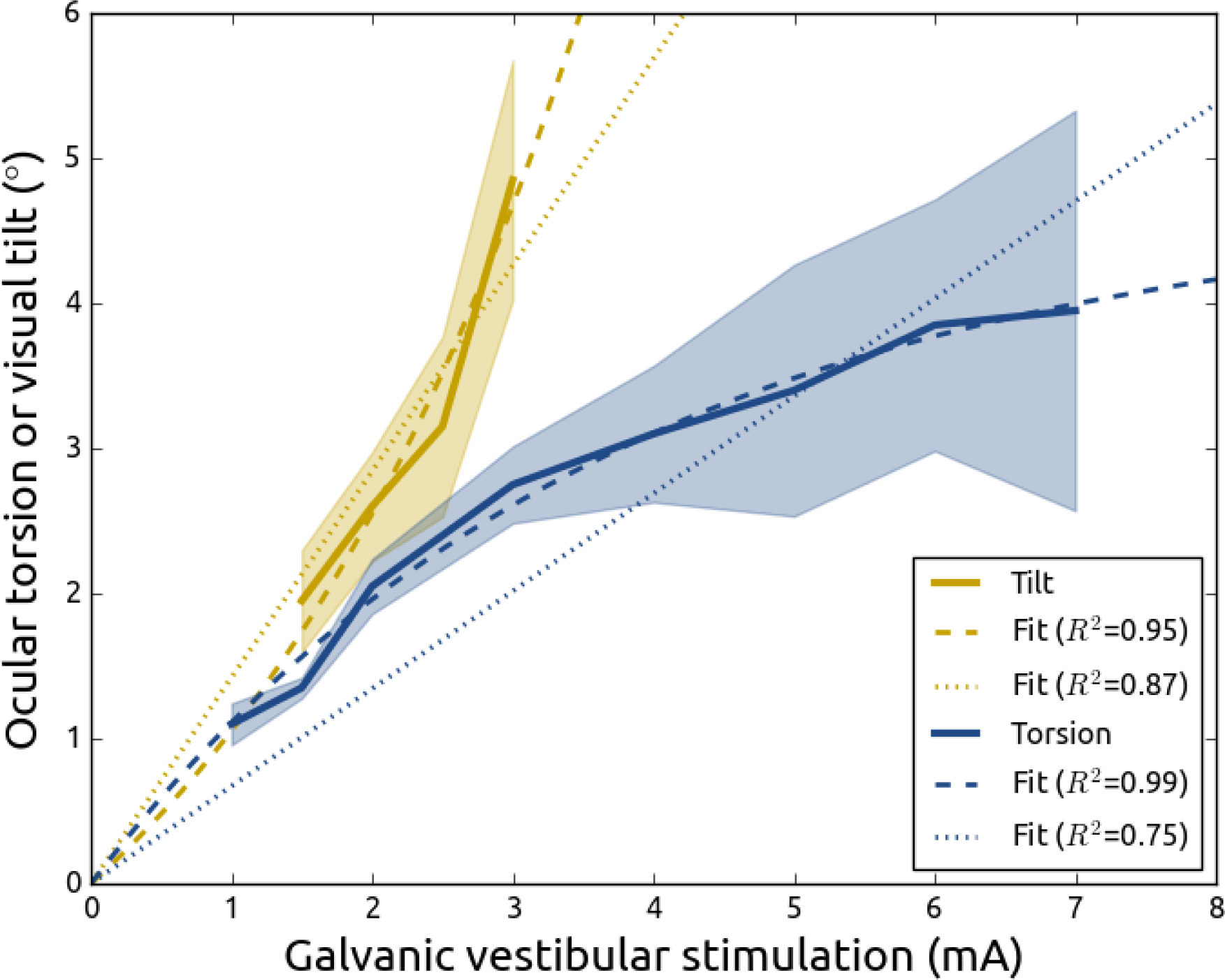
- Ocular torsion (blue) and visual tilt (yellow) in degrees (y-axis) as a function of unipolar direct current galvanic vestibular stimulation (x-axis). Solid lines represent the average and shading the standard error of the mean in data reported by Zink et al. (1998). Dotted lines represents directly proportional linear fits (Equations 3 and 4), and dashed lines represent exponential model fits (Equations 5 and 6).

A linear model with a free intercept variable (**Equation 1**) explained 94 percent of the variance, but is biologically impossible due to its baseline position of 0.913 degrees of ocular rotation at 0 mA of galvanic vestibular stimulation, and −0.913 degrees at −0 mA.

### Visual tilt

For visual tilt under galvanic vestibular stimulation (**Figure 3**, in yellow), a directly proportional model (**Equation 4**) explained 87 percent of the variance, and an exponential model (**Equation 6**) explained 95 percent.

A linear model with a free intercept variable (**Equation 2**) accounted for the same amount of variance as the directly proportional linear model, because the best fitting parameters were equal for both.

### Visual tilt as a function of ocular torsion

When visual tilt is plotted as a function of ocular torsion under the same galvanic vestibular stimulation intensities (**Figure 4**), it is best fitted by a combination of both exponential models (**Equations 5 and 6**), or by a combination of an exponential model of the relationship between galvanic vestibular stimulation and ocular torsion (**Equation 5**) and a directly proportional linear model of the relationship between galvanic vestibular stimulation and visual tilt (**Equation 4**). The relationship between ocular torsion and visual tilt under galvanic vestibular stimulation is worst fitted by a combination of directly proportional linear models of both (**Equations 3 and 4**).

**Figure 4.**
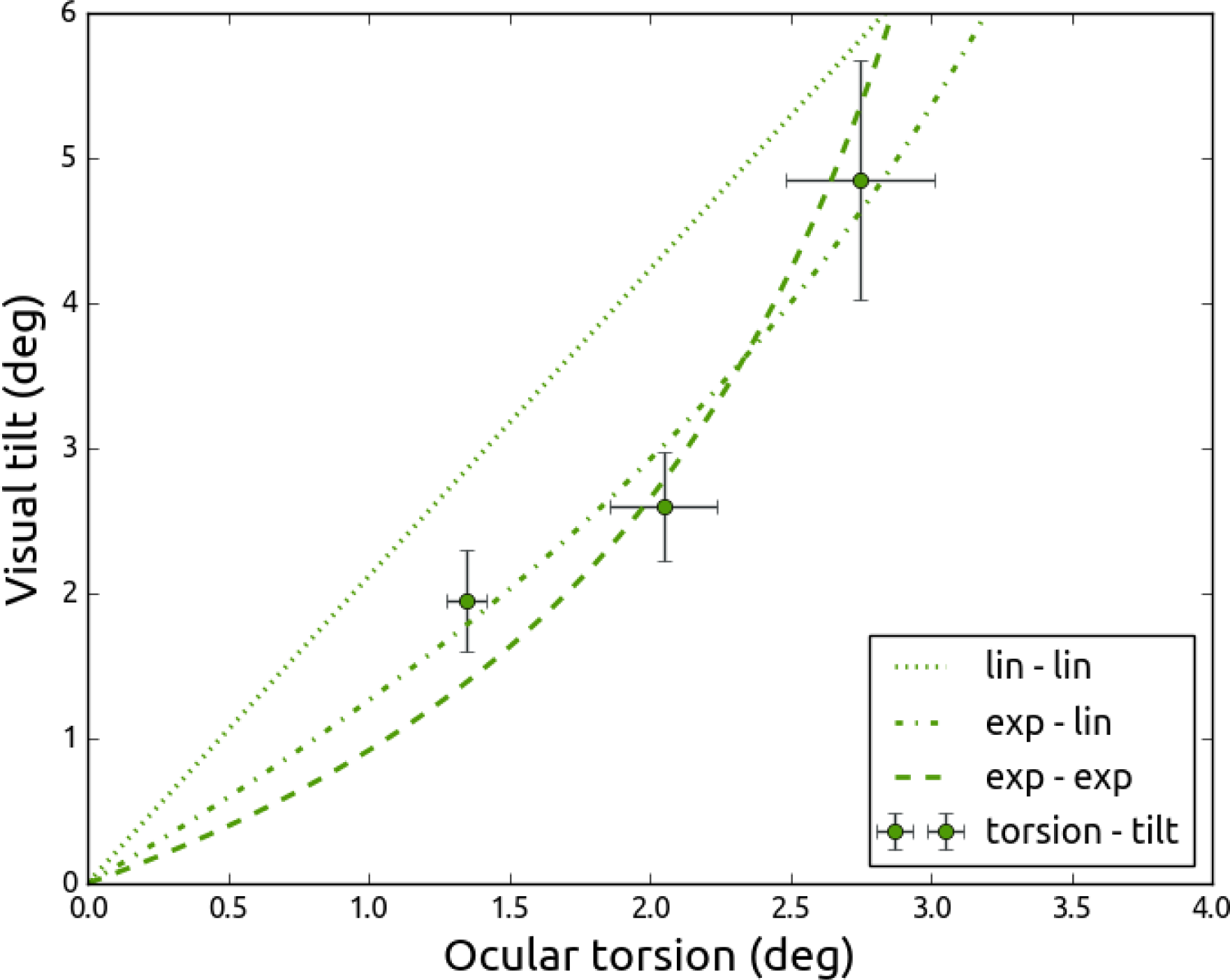
- The relationship between the effects of galvanic vestibular stimulation on ocular torsion (x-axis) and visual tilt (y-axis). Points indicate averages of data reported by Zink et al. (1998) for galvanic vestibular stimulation unipolar direct current intensities 1.5, 2.0, and 3.0 mA, and error bars indicate the standard error of the mean. The dotted line (labelled ‘lin - lin) is a combination of directly proportional linear models of the relationship between vestibular input and ocular torsion (Equation 3) or visual tilt (Equation 4). The dashed line (labelled ‘exp - exp’ is a combination of exponential models of the relationship between vestibular input and ocular torsion (Equation 5) or visual tilt (Equation 6). The dashed-dotted line (labelled ‘exp - lin’) represents a combination Equations 5 and 4. The fits are the same as those presented in Figure. 2 and 3; they are simply replotted in the same space.

## Discussion

In the current study, data from Zink and colleagues (1998, *Electroencephalography and Clinical Neurophysiology, 107)* was reanalysed using models that more accurately reflect biology (specifically the position of the eye at no vestibular input). Zink and colleagues applied galvanic vestibular stimulation, using a unipolar and direct current, and measured either ocular torsion or visual tilt. In the current study, it is demonstrated that a directly proportional linear model fitted the data from both measures relatively well. However, ocular torsion is better described by a model with an exponentially decreasing slope that moves towards an asymptote. These results show that with increasing vestibular stimulation, ocular torsion slopes down, whereas visual tilt increases (perhaps exponentially). The lack of a linear relationship between the vestibular effects on ocular torsion and on visual tilt suggests that ocular torsion is not the sole contributor of visual tilt, but that instead vestibular input could be processed centrally to maintain visual stability.

### Central processing of vestibular information

In multi-sensory research, the most simple explanation for the interaction between two senses (e.g. the vestibular and visual system) is often *stochastic resonance* (Lugo, Doti, & Faubert, 2008). Stochastic resonance occurs when the general level of activation of multi-sensory neurons is heightened by a stimulus (e.g. galvanic vestibular stimulation or auditory noise), which leads to a higher sensitivity to faint stimuli from another sense (e.g. tactile information) that on its own would lead to sub-threshold activation for detection. An example of such a study is by Ferrè and colleagues (Ferrè, Day, Bottini, & Haggard, 2013), who showed that galvanic vestibular stimulation indeed leads to a higher sensitivity for a faint tactile stimulus. They offer an explanation based in stochastic resonance, hypothesising simultaneous activation of bimodal neurons in parietal opperculum. However, the data from Zink and colleagues re-analysed here cannot be explained by stochastic resonance, as the observed visual tilt was direction-dependent on the stimulation’s current direction.

An alternative explanation is that vestibular information is processed in visual cortex. When anaesthetised cats with fixated eyeballs are tilted, the receptive fields of a proportion of neurons in visual cortex tilts as a result (Denney & Adorjanti, 1972; Horn & Hill, 1969; Horn et al., 1972). This suggests that a mechanism might exist by which head roll that is not otherwise compensated for, is compensated by the rotation of receptive fields.

Contrary to the above, non-anaesthetised cats with non-fixated eyeballs do show tilt in receptive fields in a proportion of neurons in visual cortex, but they are uncorrelated with bodily tilt (Schwartzkroin, 1972). It was suggested that receptive field tilting was a response to general arousal rather than a systematic processing of visual tilt.

Although Zink and colleagues used direct current stimulation, others have employed vestibular stimulation with a sinusoidally alternating current, and have instead argued that visual tilt can be completely accounted for by ocular torsion (Romberg, Holst, & Doden, 1951). In addition, using 100 ms pulses, Aw and colleagues found a linear relationship between current intensity and the velocity of ocular torsion (Aw, Todd, & Halmagyi, 2006). In sum, a linear relationship between vestibular input and ocular torsion does exist when pulsed or alternating current galvanic vestibular stimulation is employed.

A direct investigation of the non-linear properties of the torsional response to natural vestibular stimulation (by means of head rotation) is described by Schneider, who modelled the gain and intensity of torsional nystagmus (Schneider, Glasauer, Brandt, & Dieterich, 2003). Specifically, they demonstrate that nystagmus was present during low-frequency but not high-frequency stimulation, and argue that it is the contribution of nystagmus that causes the non-linear relationship between vestibular input and ocular torsion.

### Vestibular effects on attention

Another potentially interesting role of vestibular input is highlighted by Shuren and colleagues, who rotated participants in a revolving chair before asking them to perform a line bisection task. After leftward rotation, participants showed an increased leftward bisection error, suggesting that vestibular input might induce an attentional bias (Shuren, Hartley, & Heilman, 1998).

Further evidence for a potential role of vestibular functioning in spatial attention is provided by studies of vestibular stimulation in neglect syndrome. Neglect syndrome occurs primarily after damage to right parietal cortex, and patients display a strong attentional bias towards ipsilesional space. Neglect patients but not control patients (with similar lesions but without neglect) show deviations of their perceived visual vertical during roll head movements (Saj, Honore, Bernati, Coello, & Rousseaux, 2005), and exhibit a larger deviation of subjective visual vertical and horizontal (Kerkhoff & Zoelch, 1998). Furthermore, neglect symptoms decrease during (Rubens, 1985) or immediately after caloric vestibular stimulation, but these effects do not linger for more than 15 minutes (Cappa, Sterzi, Vallar, & Bisiach, 1987).

## Conclusion

Vestibular input by means of galvanic vestibular stimulation leads to both ocular torsion as a result of the vestibulo-ocular reflex, and visual tilt as a result of ocular torsion. However, visualt tilt is not always linearly related to ocular torsion under galvanic vestibular stimulation, hinting at an additional central processing of vestibular information to stabilise vision. One such mechanism could be the tilting of receptive fields in visual cortex, although results supporting this theory are inconsistent. A perhaps more likely explanation of the non-linear effects of vestibular input on ocular torsion comes from the observation that at low stimulation frequencies, non-linear torsional nystagmus occurs on top of slow phase ocular torsion. In conclusion, central processing of vestibular information and induced nystagmus occurs to stabilise vision during rolling head movements.

## Acknowledgements

I would like to thank Ignace Hooge, Lotte van Nierop, and Martine van Zandvoort for their comments on an earlier version of this manuscript. In addition, I am indebted to an anonymous reviewer who interpreted the presented results with a greater mastery of the literature on galvanic vestibular stimulation, in response to the submission of an earlier version of this manuscript to *Journal of Vision.*

